# Optimizing athlete assessment of maximal force and rate of development: A comparison of the isometric squat and mid-thigh pull

**DOI:** 10.1101/2020.04.17.046359

**Authors:** Joao Renato Silva, Vasileios Sideris, Bryna C.R. Chrismas, Paul J. Read

**Affiliations:** National Sports Medicine Program, Excellence in Football Project, Aspetar, Qatar Orthopedic and Sports Medicine Hospital Doha, Qatar; Center of Research, Education, Innovation and Intervention in Sports, University of Porto, 4200-450 Porto, Portugal; Department of Rehabilitation, Aspetar, Qatar Orthopedic and Sports Medicine Hospital Doha, Qatar; Qatar University, Sport Science Program, College of Arts and Science, Doha, Qatar; Athlete Health and Research Performance Center, Aspetar, Qatar Orthopedic and Sports Medicine Hospital Doha, Qatar; School of Sport and Exercise, University of Gloucestershire, Gloucester, UK

**Keywords:** isometric strength, rate of force development, unilateral and bilateral

## Abstract

This study compared force-time characteristics and muscle activity between the isometric squat (ISQ) and mid-thigh pull (IMTP) in both bilateral (ISQ_BI and_ IMTP_BI_) and unilateral (ISQ_UNI and_ IMTP_UNI_) stance. Peak force (PF), rate-of-force (RFD) (e.g. 0-300ms) and EMG of the multifidus, erector spinae (ES), gluteus maximus (GM), biceps femoris (BF), semitendinosus (ST), vastus medialis (VM), vastus lateralis (VL) and soleus were recorded in ten recreationally trained males. PF was significantly greater during the ISQ_BI_ vs. IMTP_BI_ (p=0.016, ES=1.08) but not in the unilateral test mode although effects remained moderate (ES=0.62). A trend indicated heightened RFD_300ms_ (p = 0.083; ES=0.81) during the IMTP_BI_ vs. the ISQ_BI_, but these effects were smaller in the unilateral test (ES = 0.51). Greater (p<0.0001) EMG for VL (ES=1.00-1.13) and VM were recorded during the ISQ compared to IMTP modes in both modes (ES = 0.97 – 1.18). Greater BF EMG (p = 0.030, ES = 0.31) was shown in IMTP_BI_ vs. ISQ_BI_ and these effects were stronger in the unilateral modes (p = < 0.05; ES = 0.81 – 0.83). Significantly greater ST activation was shown in both IMTP_UNI_ (p < 0.05; ES = 0.69-0.76) and IMTP_BI_ (p < 0.001; ES = 1.08). These findings indicate that ISQ results in elevated PF, whereas, RFD is heightened during the IMTP and these differences are more pronounced in bilateral modes. Greater activation of the quadriceps and hamstring muscles are expected in ISQ and IMPT respectively.

## INTRODUCTION

Muscular strength is a fundamental physical quality that underpins sports performance.^23^ Rate of force development (RFD) is also of high importance when constraints exists for the time availability of force production.^17^ Optimization of these physical characteristics through effective testing allows more targeted training interventions to be implemented.

Recently, there has been an increase in the utilization of maximal strength tests such as the isometric squat (ISQ) and mid-thigh pull (IMTP).^2,4,6,9,12^ These assessments have lower skill requirements than traditional iso-inertial testing and therefore may be more advantageous in settings such as large group programs, rehabilitation and when working with athletes of a low training age.^5^

Numerous studies have included the IMTP,^6,9,25^ with fewer employing the ISQ.^2,21^ Nuzzo et al.^21^ utilized both assessment modes and showed they were significantly correlated (r = 0.76); however, peak force and RFD were higher in the ISQ. Greater peak force during the ISQ has also been shown for females only, with a trend evident of higher impulse and RFD during the ISQ but these differences were not significant.^4^ No meaningful differences were observed between the two assessment modes in the male athletes.^4^ Cumulatively, the available evidence comparing force-time characteristics in the two tests are equivocal and require further investigation. Furthermore, previous literature investigating the differences between the ISQ and IMTP have only used a bilateral stance.^4,21^ Given the increased frequency of research pertaining to measurement of unilateral tasks,^2,3^ and their associations with reductions in performance,^3^ a more comprehensive understanding of these assessment modes is warranted.

In addition to kinetic comparisons, it is necessary to examine the level of muscle activation to further inform practitioners. This information has implications for athlete profiling, longitudinal monitoring and training prescription. In the IMTP, the arms are used to transmit forces from the ground as participants “pull” the bar, whereas in the ISQ the bar is positioned on the upper back.^4^ Although there are clearly similarities between the two actions as indicated by their significant correlation,^21^ the set-up of the IMTP may result in greater recruitment of posterior chain musculature. Conversely, the ISQ may heighten muscle activation of the knee extensors. However, this is currently speculatively and requires data to support this notion.

The purpose of this investigation was to compare the force-time characteristics and muscle activity of selected lower body and trunk muscles during both the unilateral and bilateral ISQ and IMTP. It was hypothesized that test modes would elicit different muscle activation patterns with the squat showing increased quadriceps activation and resultant peak force.

## METHODS

### Experimental design

A repeated measure, randomized, counterbalanced study design was used to control for exercise order (isometric mid-thigh pull and isometric squat respectively) and mode (unilateral and bilateral). Following a prior familiarization, all recorded tests were subsequently performed in a single testing session. Participants were required to abstain from training for 48 hours prior and asked to maintain a consistent fluid (water only) and dietary intake on the day of testing.

### Subjects

Ten recreationally trained participants (age = 37.4 ± 4.1 years; body mass = 74.4 ± 3.9 kg; height = 1.75 ± 0.04 m; body mass index 24 ±1 kg/m^2^) with no current musculoskeletal injuries, and a minimum of two years of resistance training experience volunteered to take part. This study was approved by the Anti-Doping Lab, Qatar, IRB: F2017000227 and all the subjects provided informed consent prior to participation. All procedures were conducted in accordance with the Declaration of Helsinki.

### Procedures

On arrival, height (Stadiometer: Seca, Birmingham, UK) and body mass were assessed (Seca Digital Scales, Model 707: Seca, Birmingham, UK) measured to the nearest 0.1 cm and 0.1 kg, respectively. Participants first performed a standardized 10-minute general warm-up consisting of pulse raising and dynamic stretches in addition to two practice trials at 60 and 80% of their perceived maximum effort for each exercise condition.

### Unilateral and Bilateral Isometric Squat and Mid-Thigh Pull

The IMTP and ISQ were performed in a ‘ISO rig’ (Absolute Performance, Cardiff, UK) with either one or two single axis force plates (dependent on the test mode) (PASPORT force plate, PASCO Scientific, California, USA) sampling at 1000 Hz, positioned directly underneath the steel bar. Incremental height adjustments of 1 cm were used to accommodate athletes of different statures. Set up protocol was completed in accordance with a recent commentary describing recommended test procedures.^7^ Participants were instructed to maintain an upright torso (forward lean of 5° from vertical) and a manual goniometer was used to record the knee (135° to 140°) and hip (140° to 145°) angle during both the unilateral and bilateral modes. Angles were recorded for each participant and replicated across the different test modes to allow comparison and ensure standardization. No significant differences were observed between anatomical angles irrespective of test mode (IMTP vs. ISQ) and condition (unilateral vs. bilateral) (p > 0.05). During the unilateral testing modes, the non-stance limb hovered next to the working limb to keep the hips level, aiding balance and stability. During the IMTP, participants used lifting straps to ensure grip was not the limiting factor to lower limb strength expression. Grip width and foot position were standardized within participants.^5^

Independently of exercise condition, participants were required to stand in the centre of the force plate and remain motionless for a period of at least 1s prior to the initiation of the test. Participants were instructed take the ‘slack’ from the body while being monitored to maintain minimal pre-tension (verified by manual detection of the force-time curve in real time) ensuring the force trace is similar to body mass and steady.^7^ If an uncontrolled pre-tension (>50 N change) and / or countermovement (e.g. production of a negative/antagonist force) prior the onset of the action was observed, the trial was repeated.^7^ Each trial was initiated by a “3, 2, 1, Go” countdown and participants were instructed to ‘push the ground away by driving their feet into the force plate as fast and hard as possible’.^7^ These instructions have been recommended to maximize both PF and RFD.^17^ The participants maintained the effort for 4s with strong and standardized verbal encouragement provided throughout. Two trials were performed, and the mean reported for each condition (right, left and bilateral) with 30 seconds rest between each and 2 minutes’ rest between testing modes.

Recorded metrics included net peak force (PF – body weight) and average rate of force development [(RFD, Δ Force/Δ time] at 100, 200, and 300 ms time windows. Contraction onset was determined using 5 standard deviations of body weight recorded during an initial 1s weighing period of quite standing prior to the test initiation.^8^

### Electromyography measurement

Data were collected by a sixteen-channel wireless EMG system (DTS, Noraxon USA Inc, Scottsdale, Arizona). Standard skin preparation was completed prior to dual electrode placement (Noraxon USA Inc, Scottsdale, Arizona). Self-adhesive electrodes were attached in parallel to the fibers of the multifidus, erector spinae (longissimus), gluteus maximus, biceps femoris, semitendinosus, vastus medialis, vastus lateralis and soleus on both body sides in accordance with SENIAM guidelines.^14^

Raw EMG signals were sent in real time to a computer and recorded at 1500 Hz. The data were recorded and further analyzed by MyoResearch 3.12 software (Noraxon USA Inc, Scottsdale, Arizona). Signals were high pass filtered (10 Hz), rectified and smoothed with a root mean square algorithm with a 100-millisecond window. Mean and maximum EMG data were normalized to the mean peak of a 1000-millisecond window from a 5s maximal voluntary isometric contraction (MVIC) trial for each muscle in accordance with previous guidelines.^14^ Mean and peak EMG signal amplitudes over a 3-second window after muscle onset of each condition were calculated and averaged. Muscle onset was determined as a rise at 20% of peak EMG signal of the vastus medialis muscle using MR3.12 software auto-marker algorithm and confirmed by visual inspection.

### Statistical analysis

Data are presented as mean ± standard deviations. The normality distribution of the data was checked with the Shapiro–Wilk test. To examine differences in force-time data and EMG between respective test modes (IMTP vs. ISQ) in both bilateral and unilateral stance (dominant limb), a paired samples t-test was used. Additionally, Cohen ES statistics were calculated and defined as trivial (< 0.2), small (0.2 – 0.6), moderate (0.6 – 1.2), large (1.2 – 2.0) and very large (< 2.0).^15^ Data were analyzed using Microsoft Excel and SPSS software (version 21.0, IBM SPSS Statistics, Chicago, IL, USA).

## RESULTS

### Force Production

PF and RFD measures during the ISQ and IMTP tests in the unilateral and bilateral modes are presented in Table 1. Data revealed significantly greater PF during the ISQ_BI_ (p = 0.016) only. Moderate effect sizes were shown indicating increased PF for ISQ_UNI_ (ES = 0.62) and ISQ_BI_ (ES = 1.08) respectively. There was a trend to suggest greater RFD300ms during the IMTP_BI_ vs. ISQ_BI_ (p = 0.083) with small (ES = 0.51) and moderate effects (ES = 0.81) reported in the unilateral and bilateral mode comparison respectively. Moderate effects (ES = 0.63) were also shown for the IMTP_BI_ vs. ISQ_BI_ assessment of RFD200ms while all other comparisons were non-significant and displayed small effects (< 0.6).

**Table 1.** Peak Force and rate of force development during the different exercise modes (mean ± SD)

|  | Unilateral (n = 18) |  |  | Bilateral (n = 9) |  |  |
| --- | --- | --- | --- | --- | --- | --- |
| | IMTP<sub>UNI</sub> | ISQ<sub>UNI</sub> | $\Delta$ | IMTP<sub>BI</sub> | ISQ<sub>BI</sub> | $\Delta$ |
| **Peak Force** | 1456 (189) | 1612(284) * | 10.7% | 1771 (171) | 2111 (310) * | 19.2% |
| **RFD 100 ms** | 2730 (1091) | 2755 (1265) | 0.9% | 4364 (1882) | 3450 (13851) | -20.9% |
| **RFD 200 ms** | 2971 (980) | 2814 (847) | -5.3% | 4358 (1174) | 3651 (1221) | -16.2% |
| **RFD 300 ms** | 2888 (854) | 2555 (724) | -11.5% | 3935 (1012) | 3151 (950) | -19.9% |

RFD-rate of force development, IMTP - isometric mid-thigh pull; ISQ - isometric squat; Uni-unilateral; BI - bilateral \* $<0.05$ significant difference from IMTP

### Muscle Activity

Muscle activation during the ISQ and IMTP exercises in the unilateral and bilateral modes are displayed in Table 2. Greater peak and mean EMG activity of the vastus lateralis and medialis (p < 0.001) were present during the ISQ_UNI_ and ISQ_BI_ compared to both IMTP modes, corresponding to a moderate effect (ES = 0.97 – 1.18). There was a trend for greater peak EMG activity of the Biceps femoris (p = 0.051; ES = 0.27) in the IMTP_BI_ compared to the ISQ_BI_ and these differences were significant, but effects were small for mean EMG (p = 0.030, ES = 0.31). This observed trend became more apparent in the unilateral modes, with significant between test differences indicating the IMTP_UNI_ resulted in significantly greater peak and mean EMG biceps femoris activity corresponding to a moderate effects size (p = < 0.05; ES = 0.81 – 0.83). When observing the EMG of the semitendinosus muscle, there was a stronger effect of exercise condition, whereby significantly greater peak and mean muscle activation was shown in both IMTP_UNI_ (p < 0.05; ES = 0.69-0.76) and IMTP_BI_ (p < 0.001; ES = 1.08) in comparison to both ISQ modes. No meaningful differences or substantial effects were indicated between testing modes for any of the other muscles tested.

**Table 2.** Surface electromyography data recorded during the different exercise modes (mean ± SD)

|  | PEAK EMG (%MVIC) |  |  | MEAN EMG (%MVIC) |  |  | PEAK EMG (%MVIC) |  |  | MEAN EMG (%MVIC) |  |  |
| --- | --- | --- | --- | --- | --- | --- | --- | --- | --- | --- | --- | --- |
| | IMTP<sub>UNI</sub> | ISQ<sub>UNI</sub> | $\Delta$ | IMTP<sub>UNI</sub> | ISQ<sub>UNI</sub> | $\Delta$ | IMTP<sub>BI</sub> | ISQ<sub>BI</sub> | $\Delta$ | IMTP<sub>BI</sub> | ISQ<sub>BI</sub> | $\Delta$ |
| Erector Spinae | 83.95 (22.3) | 80.1 (36.3) | -2.1% | 40.7 (12.4) | 39.1 (24.8) | -0.5% | 113.8 (32.5) | 105.0 (34.4) | -7.9% | 71.1 (18.7) | **60.3 (19.8)\*** | -8.6% |
| Multifidus | 95.9 (32.9) | 93.9 (42.9) | -4.5% | 55.9 (20.1) | 55.6 (29.7) | -3.9% | 114.1 (31.7) | 105.6 (36.6) | -7.1% | 72.9 (19.8) | 66.6 (27.5) | -15.2% |
| Gluteus Maximus | 98.2 (41.0) | 100.6 (74.1) | 2.4% | 57.9 (25.3) | 60.3 (47.4) | 4.1% | 43.7 (19.8) | 41.9 (26.3) | -4.1% | 23.3 (9.2) | 24.3 (16.4) | 4.3% |
| Biceps Femoris | 121.8 (48.8) | **85.4 (31.7)\*** | -29.9% | **76.5 (32.8)** | **51.3 (21.6)\*** | -32.9% | 46.4 (33.0) | 37.2 (22.5) | -19.8% | 25.4 (20.5) | 20.1 (15.5) | -20.9% |
| Semitendinosus | 74.4 (18.4) | 60.1 (36.1) | -19.2% | **44.2 (11.2)** | **34.3 (22.4)\*** | -22.4% | 51.3 (14.0) | **34.7 (10.6)\*\*** | -32.4% | 31.1 (10.2) | **18.8 (7.9)\*\*** | -39.5% |
| Vastus Lateralis | 120.0 (41.0) | **182.9 (65.1)\*\*** | 52.4% | **69.1 (24.5)** | **114.3 (40.7)\*\*** | 65.0% | 74.1 (45.2) | **115.0 (39.6)\*\*** | 55.2% | 43.0 (24.6) | **72.5 (24.9)\*\*** | 68.6% |
| Vastus Medialis | 116.6 (33.0) | **192.2 (55.1)\*\*** | 64.8% | **76.4 (51.3)** | **133.4 (67.0)\*\*** | 74.1% | 83.5 (80.2) | **136.0 (61.5)\*\*** | 62.9% | 43.4 (36.7) | **81.5 (36.0)\*\*** | 87.8% |
| Soleus | 189.1 (141.4) | 196.1 (82.5) | 3.7% | 92.3 (54.2) | 110.5 (28.3) | 19.2% | 123.0 (71.4) | 156.3 (64.7) | 26.8% | 54.3 (34.4) | 77.7 (33.8) | 43.1% |

EMG- electromyography; MVIC- maximal voluntary isometric contraction; IMTP - isometric mid-thigh pull; ISQ - isometric squat;
Uni- unilateral; BI - bilateral
\*<0.05 significant difference from IMTP
\*\*<0.01 significant difference from IMTP

## DISCUSSION

The purpose of this investigation was to compare the force-time characteristics and muscle activity of selected lower body and trunk muscles during both the unilateral and bilateral ISQ and IMTP. Greater peak force values were shown during the ISQ versus the IMTP. Conversely, there was a trend for a greater rate of force development (RFD_300ms_) during the IMTP, with these differences more pronounced when comparing the bilateral exercise modes. Additionally, greater EMG activity of the anterior (vastus lateralis and medialis) and posterior (semitendinosus and bicep’s femoris) muscles of the thigh were indicated for the ISQ and IMTP respectively.

The greater sensitivity of the bilateral testing modes to differentiate the force-time characteristics specific to each task may be attributed to the decrease in base of support that occurs during unilateral tasks.^11^ The increased instability present during single leg stance may effect technique execution, decreasing maximum and rapid force generation.^11^ Several investigations have also shown that maximal isometric strength assessments used to calculate RFD are susceptible to enhanced measurement error,^13^ with shorter time-bands appearing to be less reliable for RFD assessment.^4,5^ This lower performance reproducibility could be more evident during unilateral assessments and may in part explain the clearer differences (effect sizes) in RFD observed between the ISQ and IMTP during the bilateral tasks. Cumulatively, this study indicates that force-time characteristics are different between tasks (IMTP vs. ISQ) and to optimize the measurement of maximal force-time characteristics, bilateral tasks may be more advantageous.

Our results confirm recent observations of greater peak force expression during the ISQ.^4,21^ Greater peak forces during ISQ vs. IMTP may be explained at least in part by differences in cross sectional area between the quadriceps vs hamstrings muscles groups.^22,26^ Previous research has indicated that independent of the testing mode (isometric, concentric and eccentric), higher peak forces are reported when measuring force capabilities of quadriceps vs. hamstrings muscles.^24^ In the current study, heightened recruitment of the quadriceps was shown during the ISQ and this may have resulted in a greater degree of force/tension during the ISQ independently of the testing mode.^22,26^

An elevated instability component may also affect the magnitude of muscle activation. Heightened recruitment of the quadriceps and hamstring musculature was indicated in both the ISQ_UNI_ and IMTP_UNI_ tasks. There are contradictory reports regarding greater motor unit activation during unilateral vs. bilateral maximal muscle voluntary contractions.^11,19^ Some studies have shown that the concurrent request to stabilize and maximize force production during more unstable exercises may result in decreased activation when compared to those with lower stabilization requirements.^18^ Nevertheless, this has not been observed in other studies.^11,19,20^ Several methodological factors may explain the distinct observations between studies, including ours. These include different loading schemes (maximal voluntary contractions vs. 3RM vs. unweighted), muscle actions applied (e.g. dynamic vs isometric muscle actions) and exercise modes (single vs multi-joint).^11,18,19,20^

The heightened quadriceps and hamstring recruitment shown during the ISQ and IMTP respectively could also in part be attributed to bar position as this is likely a determinant moderator of the loading requirements in each individual exercise,^16^ influencing the kinetic, kinematic and muscle activation patterns.^10^ Placing the moment arm further in front of the hip (IMTP) or behind the knee (ISQ) will distinctly affect the muscle recruitment strategy and may explain the observed changes in posterior and anterior muscular recruitment during the IMTP and ISQ respectively.^16^

When interpreting the data from the current study, some limitations should be acknowledged. A small sample size was recruited, and the analysis may have been under powered. However, additional statistical analysis was included and effect sizes to provide a clearer interpretation of the data and the strength of any effects identified. In addition, the population tested were resistance trained individuals and not team sport athletes; thus, it is difficult to generalize the findings to the larger population. We included those with substantial resistance training experience to ensure that differences identified between the tests were not based on un-familiarity. However, further research is required in other cohorts to examine if the results are replicated.

## CONCLUSIONS

In summary, the results of the current study indicate that unilateral and bilateral ISQ results in greater absolute peak force values, whereas, RFD is heightened during the IMTP. The bilateral exercise modes seem to further optimize the assessment of these the distinct force-time characteristics. The muscle activity may also vary between tasks with the ISQ resulting in a greater activation of the quadriceps muscles and the IMTP of the hamstring muscles.

Using these data, assessments can be tailored according to individual needs of the athlete based on the sport demands and if maximal strength is monitored during injury rehabilitation. The data indicate that bilateral variations are preferred to measure an individual’s true maximum strength (ISQ) and RFD capacity (IMTP). In addition, the greater quadriceps activation during the ISQ and higher peak forces may have connotations for athletes where quadriceps strength may be deficient; such as, following anterior cruciate ligament reconstruction.

## REFERENCES

1. Behm D, Anderson K, Curnew S. Muscle force and neuromuscular activation under stable and unstable conditions. J Strength Cond Res. 16: 416–422, 2002.

2. Bishop C, Lake J, Loturco I, Papadopoulos K, Turner A, Read P. Interlimb assymetries: The need for an individual approach to data analysis. J Strength Cond Res. Published ahead of print. 2018. doi: 10.1519/JSC.0000000000002729.

3. Bishop C, Turner A, Read P. Effects of inter-limb asymmetries on physical and sports performance: a systematic review. J Sports Sci. 36: 1135–1144, 2018.

4. Brady C, Harrison AJ, Flanagan E, Haff GG, Comynsm T. A Comparison of the Isometric Mid-Thigh Pull and Isometric Squat: Intraday Reliability, Usefulness and the Magnitude of Difference Between Tests. Int J Sports Physiol Perform. 13: 844–852, 2017.

5. Brady C, Harrison AJ, Comyns TM. a review of the reliability of biomechanical variables produced during the isometric midthigh pull and isometric squat and the reporting of normative data. Sports Biomech. 19: 1–25, 2020.

6. Comfort P, Jones P, McHahon J, Newton R. Effect of Knee and Trunk Angle on Kinetic Variables During the Isometric Mid-Thigh Pull: Test-Retest Reliability. Int J Sports Physiol Perform. 10: 58–63, 2015.

7. Comfort P, Dos Santos T, Beckham G, Stone MH, Guppy S, Haff G. Standardization and methodological considerations for the isometric midthigh pull. Strength Cond J 41: 57–79, 2019.

8. Dos Santos T, Jones PA, Comfort P, Thomas C. Effect of different onset thresholds on isometric midthigh pull force-time variables. J Strength Cond Res 31: 3463–3473, 2016.

9. Dos’Santos T, Thomas C, Comfort P, McMahon JJ, Jones PA. Relationships between Isometric Force-Time Characteristics and Dynamic Performance. Sports (Basel). 5, 68, 2017.

10. Ebben WP, Feldmann A, Dayne D, Mitsche P, Alexander K, Knetzger KJ. Muscle activation during lower body resistance training. Int J Sports Med. 30: 1–8, 2009.

11. Eliassem W, Saeterbakken AH, Van den Tillaar R. Comparison of bilateral and unilateral squat exercises on barbell kinematics and muscle activation. The International Journal of Sports Physical Therapy. 13: 871–881, 2018.

12. Haff GG, Stone M, O’Bryant HS, Harman E, Dinan C, Johnson R, Han K. Force-Time Dependent Characteristics of Dynamic and Isometric Muscle Actions. J Strength Cond Res. 11: 269–272, 1997.

13. Hart NH, Nimphius S, Cochrane JL, Newton RU. Reliability and validity of unilateral and bilateral isometric strength measures using a customized portable apparatus. Journal of Australian Strength & Conditioning. 20(Supplement 1): 61–67, 2012.

14. Hermens HJ, Freriks B, Disselhorst-Klug C, Rau G. Development of recommendations for SEMG sensors and sensor placement procedures. J Electromyogr Kinesiol. 10: 361–74, 2000.

15. Hopkins WG, Marshall SW, Batterham AM, Hanin J. Progressive statistics for studies in sports medicine and exercise science. Med Sci Sports Exerc. 41: 3–13, 2009.

16. Keogh J, Lake JP, Swinton PA. Practical applications of biomechanical principles in resistance training: moments and moments arms. Journal of fitness research. 2: 39–48, 2013.

17. Maffiuletti NA, Aagaard P, Blazevich AJ, Folland J, Tillin N, Duchateau J. Rate of force development: physiological and methodological considerations. Eur J Appl Physiol. 116: 1091–116, 2016.

18. McBride J, Cormie P, Deane R. Isometric squat force output and muscle activity in stable and unstable conditions. J Strength Cond Res. 20: 915–918, 2006.

19. McCurdy K, O’Kelley E, Kutz M, Langford G, Ernest J, Torres M. Comparison of Lower Extremity EMG Between the 2-Leg Squat and Modified Single-Leg Squat in Female Athletes. Journal of Sport Rehabilitation. 19: 57–70, 2010.

20. McCurdy K, Kutz M, O’Kelley E, Langford G, Ernest J. External oblique activity during the unilateral and bilateral fre weight squat. Clin Kinesiol. 64: 16–21, 2010.

21. Nuzzo JL, McBride J, Cormie P, McCaulley M. Relationhip between coutermovement jump performance and multijoint isometric and dynamic test of strength. J Strength Cond Res. 22: 699–707, 2008.

22. Overend TJ, Cunningham DA, Kramer JF, Lefcoe MS, Paterson DH. Knee extensors and knee flexors strength: Cross-sectional area ratios in young and elderly men. Journal of Gerontology: Medical Sciences. 47: 204–210, 1992.

23. Suchomel T, Nimphius S, Stone M. The Importance of Muscular Strength in Athletic Performance. Sports Med. 46: 1419–1449, 2016.

24. Silva JR, Magalhaes J, Ascensao A, Seabra AF, Rebelo AN. Training status and match activity of professional soccer players throughout a season. J Strength Cond Res. 27: 20–30, 2013.

25. Thomas C, Dos’Santos T, Comfort P, Jones PA. Relationships between unilateral muscle strength qualities nd change of direction in adolescent team-sports atlhetes. Sports (Basel). 6(3), 2018.

26. Wickiewicz TL, Roy RR, Powell PL, Edgerton VR. Muscle architecture of the human lower limb. Clin Orthop Relat Res. 179: 275–83, 1983.

